# Starvation induces extracellular accumulation of polyphosphate in *Dictyostelium discoideum* to inhibit macropinocytosis, phagocytosis, and exocytosis

**DOI:** 10.1101/2023.02.16.528874

**Authors:** Ramesh Rijal, Issam Ismail, Shiyu Jing, Richard H. Gomer

## Abstract

*Dictyostelium discoideum* is a soil-dwelling unicellular eukaryote that accumulates extracellular polyphosphate (polyP). At high cell densities, when the cells are about to overgrow their food supply and starve, the corresponding high extracellular concentrations of polyP allow the cells to preemptively anticipate starvation, inhibit proliferation, and prime themselves to begin development. In this report, we show that starved *D. discoideum* cells accumulate cell surface and extracellular polyP. Starvation reduces macropinocytosis, exocytosis, and phagocytosis, and we find that these effects require the G protein-coupled polyP receptor (GrlD) and two enzymes, Polyphosphate kinase 1 (Ppk1), which is required for synthesizing intracellular polyP, cell surface polyP, and some of the extracellular polyP, and Inositol hexakisphosphate kinase (I6kA), which is required for cell surface polyP and polyP binding to cells, and some of the extracellular polyP. PolyP reduces membrane fluidity, and we find that starvation reduces membrane fluidity, and this effect requires GrlD and Ppk1 but not I6kA. Together, these data suggest that in starved cells, extracellular polyP decreases membrane fluidity, possibly as a protective measure. In the starved cells, sensing polyP appears to decrease energy expenditure from ingestion, and decrease exocytosis, to both decrease energy expenditures and retain nutrients.

## Introduction

Eukaryotic cells possess the ability to uptake particles by phagocytosis or fluid by macropinocytosis. Phagocytosis and macropinocytosis share a common evolutionary origin, and these processes co-evolved as feeding mechanisms [1]. *Dictyostelium discoideum* is a soil amoeba that feeds on bacteria by phagocytosis [1–3]. Some *D. discoideum* strains can be grown axenically in a defined liquid nutrient medium, and these axenic strains possess an increased rate of macropinocytosis [4–9]. The axenic strains have a mutation in the gene encoding the Ras GTPase activating protein Neurofibromin (NF1) that regulates both phagocytosis and macropinocytosis [10], and loss of NF1 potentiates Ras activation at the sites where membrane ruffles form during macropinocytosis, thus increasing macropinocytosis [10]. Following endocytosis, the particle or fluid is transferred to lysosomeswhere the particle or fluid is digested, and the undigested material is then exocytosed [11, 12].

In nutrient-rich conditions, *D. discoideum* exists as unicellular amoebae. Starvation initiates a developmental cycle where cells aggregate together to form a multicellular fruiting body consisting of a mass of spores held off the substrate by a column of stalk cells [13, 14]. During development, starting 1 hour after starvation, *D. discoideum* cells suppress macropinocytosis by ~80% [15], and after 6 hours of starvation, *D. discoideum* cells suppress phagocytosis by ~50% [16]. At 8 hours of starvation, most *D. discoideum* cells have almost completelyreduced both macropinocytosis and phagocytosis [16]. However, during development, a small percentage of cells maintain high levels of phagocytosis and behave like immune cells that patrol the multicellular structures [17, 18]. In addition, starting 1 hour after starvation, *D. discoideum* cells reduce exocytosis, and at 4 hours exocytosis is reduced by > 50% [15]. How *D. discoideum* cells inhibit macropinocytosis, phagocytosis, and exocytosis during development remains unclear.

Polyphosphate (polyP) is a polymer of inorganic phosphate residues linked by phosphoanhydride bonds. PolyP is present in all life kingdoms [19], and it is involved in energy and phosphate storage, survival under stress conditions, and biofilm formation, and virulence in prokaryotes, and blood coagulation, inflammation, proliferation of leukemia cells, and bone calcification in eukaryotes [19–26]. Proliferating *D. discoideum* cells continuously secrete polyP, and when the cells have reached a high cell density where the cells will be about to overgrow their nutrient source, the corresponding high extracellular polyP concentrations inhibit proliferation, reduce phagocytosis, macropinocytosis, and exocytosis, and induce aggregation [24, 27, 28]. *D. discoideum* synthesizes polyP using polyphosphate kinase 1 (Ppk1), a highly conserved enzyme in prokaryotes, and likely to have been acquired by *D. discoideum* through horizontal gene transfer [29, 30]. *D. discoideum* cells lacking Ppk1 have undetectable levels of intracellular polyP [30], and reduced but detectable levels of extracellular polyP [27]. PolyP levels are also reduced in cells lacking inositol hexakisphosphate kinase A (I6kA), an enzyme that phosphorylates the inositol pyrophosphate IP6 to generate IP7 [27, 31]. PolyP shows saturable binding to *D. discoideum*, and the binding is lost in cells lacking the G protein-coupled receptor GrlD [27]. GrlD is required for polyP inhibition of proliferation [32]. In addition, GrlD, Ppk1, and I6kA are all required for polyP inhibition of macropinocytosis and exocytosis in proliferating cells [27, 28]. Although polyP inhibits phagocytosis in proliferating cells [28], we do not know whether GrlD, Ppk1, or I6kA are required for this inhibition.

PolyP induces aggregation of *D. discoideum* cells even in the presence of nutrients [24], suggesting that high levels of extracellular polyP may act as a signal that lets cells sense that they are at a high cell density, and thus probably about to overgrow their food supply and starve. This in turn would allow cells to be able to anticipate starvation and begin development. Throughout development, *D. discoideum* cells accumulate high concentrations of intracellular polyP, and the concentrations reach > 100-fold in the fruiting body compared to vegetative cells [30, 33],

Little is known about the function of polyP during *D. discoideum* development. In this report, we examined how *D. discoideum* regulates polyP production in response to starvation. We found that after 6 hours of starvation, *D. discoideum* accumulates large amounts of secreted and cell-surface extracellular polyP, and the development-associated reduction in macropinocytosis, exocytosis, and phagocytosis requires GrlD, Ppk1, and I6kA. This then suggests that the development-associated reduction in macropinocytosis, exocytosis, and phagocytosis is caused by high extracellular levels of polyP sensed by the receptor GrlD.

## Results

### Starving *D. discoideum* cells require GrlD and Ppk1 to accumulate normal levels of extracellular and cell surface polyP, and I6kA to accumulate normal levels of cell surface polyP

Proliferating *D. discoideum* cells accumulate extracellular polyP, and loss of either Ppk1 or I6kA reduce this accumulation by ~ 50%, although only at very high cell densities [27]. The cells then require GrlD to sense the extracellular polyP [32]. Starvation induces the accumulation of intracellular polyP in *D. discoideum* cells [30, 33]. To determine if *D. discoideum* cells accumulate extracellular polyP during starvation, wild-type (WT) cells or cells lacking GrlD (*grlD*^−^), polyphosphate kinase 1 (*ppk1*^−^), or inositol hexakisphosphate kinase A (*i6kA*^−^) were incubated for 6 hours in the defined growth medium SIH, or the starvation buffer PBM. Conditioned medium (CM) was collected, and polyP was measured using DAPI. As previously observed [27], mid-log phase WT, *grlD*^−^, *ppk1*^−^, and *i6kA*^−^ cells had comparable levels of extracellular polyP (**Fig. 1A**). Compared to cells in SIH or cells starved for 2 hours (**Fig. 1A and B**), extracellular polyP levels in WT cells increased after 6 hours in PBM (**Fig. 1B**). Compared to WT cells, *grlD*^−^ and *ppk1*^−^ cells accumulated less extracellular polyP after starvation for 6 hours (**Fig. 1B**). The *i6kA*^−^ cells had extracellular polyP concentrations similar to WT (**Fig. 1A and B**). Together, these data suggest that *D. discoideum* cells accumulate extracellular polyP during the early stages of starvation, that I6kA is not necessary for this effect, and that Ppk1 mediates some but not all of the extracellular polyP accumulation. Possibly because sensing polyP is needed for cells to accumulate polyP, cells also require GrlD to increase extracellular polyP at 6 hours of starvation.

**Figure 1.**
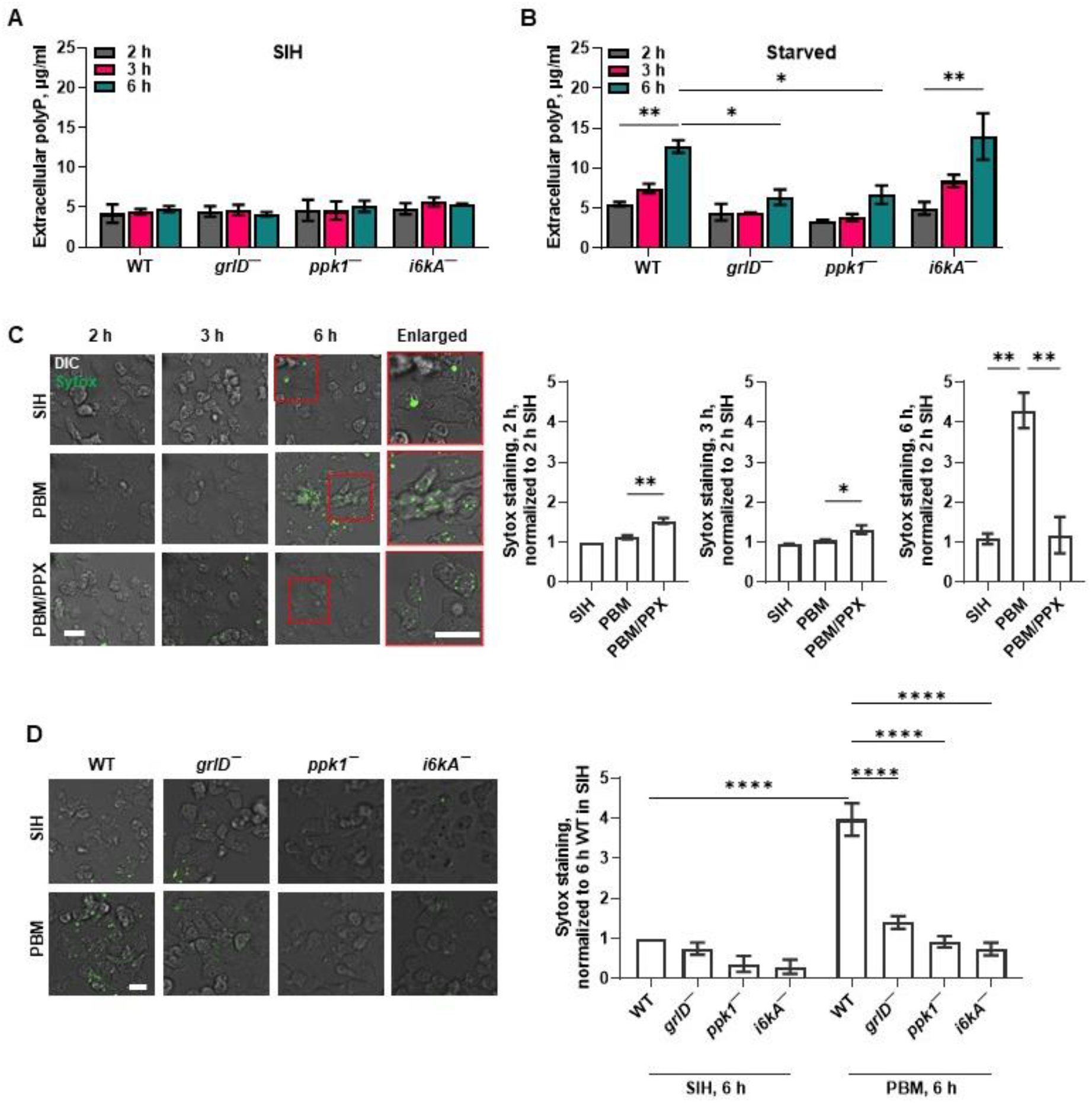
Starving *D. discoideum* cells require GrlD and Ppk1 to accumulate extracellular polyP, and GrlD, Ppk1, and I6kA to accumulate cell surface polyP. **A-B)** Wild-type (WT), *grlD*^−^, *ppk1*^−^, and *i6kA*^−^ cells were incubated in SIH (A) or starved in PBM (B) for 2, 3, and 6 hours (h) and extracellular polyP concentrations were measured at the indicated times. **C)** WT cells were incubated in SIH, PBM, or PBM with PPX for 2, 3, and 6 h, stained with SYTOX (green), and the mean fluorescence intensities of SYTOX staining were measured at the indicated times. The mean fluorescence intensity of SYTOX staining in cells incubated in SIH for 2 h was considered 1. Representative images of 3 independent experiments are shown. Cell boundaries in the enlarged insets show the SYTOX staining (green) of cells. DIC indicates differential interference contrast. Bars are 10 μm. **D)** WT, *grlD*^−^, *ppk1*^−^, and *i6kA*^−^ cells were incubated in SIH or PBM for 6 h, stained with SYTOX, and the mean fluorescence intensities of SYTOX staining were measured. The mean fluorescence intensity of SYTOX staining in cells incubated in SIH for 6 h was considered 1. Representative images of 3 independent experiments are shown. Bar is 10 μm. Values are mean ± SEM of 3 independent experiments (A - D). * indicates p < 0.05, ** p < 0.01, and **** p < 0.0001 (One-way ANOVA with Dunnett’s multiple comparisons test (B and C)) (Two-way ANOVA with Bonferroni’s multiple comparisons test (D)).

PolyP forms condensed spherical nanoparticles on the surface of activated human platelets [34]. Cell-surface-associated polyP can be detected with the cell-impermeable high affinity nucleic acid stain SYTOX [34]. To determine if extracellular polyP is present on the cell surface, cells were stained with SYTOX. We found that WT cells starved for 6 hours showed increased SYTOX staining compared to cells in SIH (**Fig. 1C and D**). To determine if the SYTOX staining was due to nucleic acid that is present on the cell surface, for instance due to release from dead cells, WT cells were starved in PBM for 5.5 hours, then ribonuclease (RNase) or deoxyribonuclease (DNase) were added to cells, and the cells were incubated for 30 minutes, and then SYTOX staining of the cells was performed. We also tested the enzymatic activity of RNase and DNase by incubating RNA or DNA with RNase or DNase at the concentration that was used to treat the cells. At concentrations where the RNase or DNase completely digested RNA or DNA, respectively (**Fig. 2A**), RNase or DNase did not significantly affect the fluorescence intensity of the SYTOX staining (**Fig. 2B**), suggesting that the fluorescence on cells from SYTOX staining is not due to the presence of extraneous nucleic acids but likely due to the presence of polyP.

**Figure 2.**
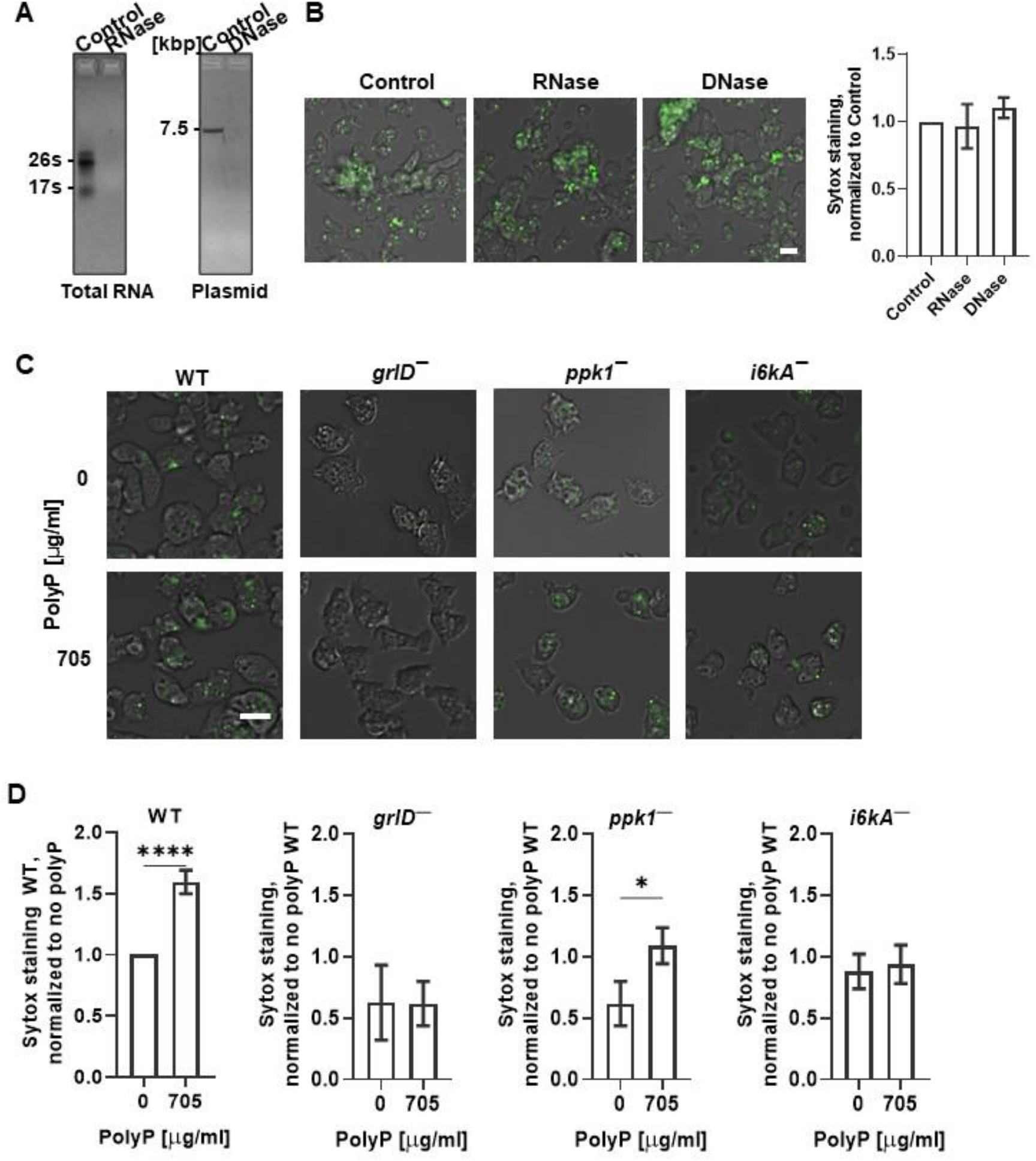
Starving *D. discoideum* cells require GrlD and I6kA to bind exogenous polyP. **A)** Agarose gels of total RNA or plasmid DNA untreated (Control) or treated with RNase or DNase, respectively, stained with ethidium bromide are shown. Kbp indicates kilo base pair, and s indicates Svedbergs, a unit for sedimentation coefficient. **B)** WT cells starved in PBM for 6 hours were treated with RNase or DNase and stained with SYTOX, and the normalized mean fluorescence intensities of SYTOX staining were determined. **C)** Vegetative WT, *grlD*^−^, *ppk1*^−^, and *i6kA*^−^ cells were incubated no polyP (0) or with 705 μg/ml polyP and stained with SYOTX. For B and C, representative merged images of differential interference contrast (DIC) and the SYTOX staining (green) are shown from 3 independent experiments. Bars are 10 μm. **D)** Quantification of the mean fluorescence intensities of SYTOX staining from C are shown. The mean fluorescence intensity of the SYTOX staining in WT cells with no polyP (0) was considered 1. Values in B and D are mean ± SEM of 3 independent experiments. * indicates p < 0.05 and **** p < 0.0001 (t tests).

In the control assays for Fig. 1C and D, cells were incubated in PBM in the presence of 5 μg/ml of yeast exopolyphosphatase (PPX), an enzyme that degrades polyP by removing terminal phosphate residues [35]. At 2 and 3 hours of incubation, the SYTOX staining was higher in the PPX-treated group than in the cells incubated in SIH or PBM alone. It is possible that *D. discoideum* might compensate for an initial loss of polyP when exposed to PPX by synthesizing more polyP. However, in cells incubated in PBM with PPX at 6 hours, the fluorescence intensity of the SYTOX staining was significantly lower than in the cells incubated in SIH or PBM alone (**Fig. 1C**). This suggests that cells do not compensate for a persistent degradation of extracellular polyp or cannot maintain high levels of production, and that the SYTOX staining at 6 hours of starvation may be due to polyP on the cells. In SIH, *ppk1*^−^ and *i6kA*^−^ appeared to have decreased SYTOX staining, but this was not statistically significant (**Fig. 1D**). For cells starved for 6 hours, compared to WT cells, *grlD*^−^, *ppk1^−^*, and *i6kA*^−^ cells showed decreased SYTOX staining (**Fig. 1D**). These data suggest that in addition to the accumulation of extracellular polyP at 6 hours of starvation, *D. discoideum* cells accumulate cell surface polyP at 6 hours of starvation, and that this process is potentiated by GrlD, Ppk1, and I6kA.

### Cells require GrlD and I6kA to bind exogenous polyP

WT *D. discoideum* cells bind extracellular polyP, and the loss of GrlD reduces the binding of polyP to cells [27, 32]. To test if the binding of extracellular polyP to WT cells increases the SYTOX staining, cells in SIH were incubated with exogenous polyP and incubated with SYTOX. As in Figure 1D, in SIH in the absence of exogenous polyP, WT, *grlD*^−^, *ppk1*^−^, and *i6kA*^−^ cells showed similar levels of SYTOX staining. Incubation with exogenous polyP increased the SYTOX staining of WT and *ppk1*^−^ cells, but not *grlD*^−^ or *i6kA*^−^ cells (**Fig. 2C and D**). Together, the data suggest that SYTOX shows a basal staining on cells, and that GrlD and I6kA are needed for cells to bind additional exogenous polyP.

### Starvation reduces the cell membrane fluidity of *D. discoideum* cells, and this requires GrlD and Ppk1

Cell membrane physical properties such as membrane fluidity are important regulators of endocytosis and exocytosis in mammalian cells [36, 37]. We previously observed that in SIH, WT, *grlD*^−^, *ppk1*^−^, and *i6kA*^−^ cells have similar membrane fluidity, that polyP decreases membrane fluidity of WT cells, and that this requires GrlD, Ppk1, and I6kA [28]. Compared to WT and *i6kA*^−^ cells in SIH [28] or freshly starved WT cells, WT and *i6kA*^−^ cells starved for 6 hours showed a decreased membrane fluidity as indicated by an increased half-life of recovery, decreased diffusion coefficient, and decreased mobile fraction (**Fig. 3A and B**). Compared to cells in SIH ([28] and **Fig. 3B**), the 6-hour starved *grlD*^−^ and *ppk1*^−^ cells did not show decreased membrane fluidity (**Fig. 3B**). Together, these data suggest that starvation reduces cell membrane fluidity in *D. discoideum* cells, and that this effect requires GrlD and Ppk1 but not I6kA.

**Figure 3.**
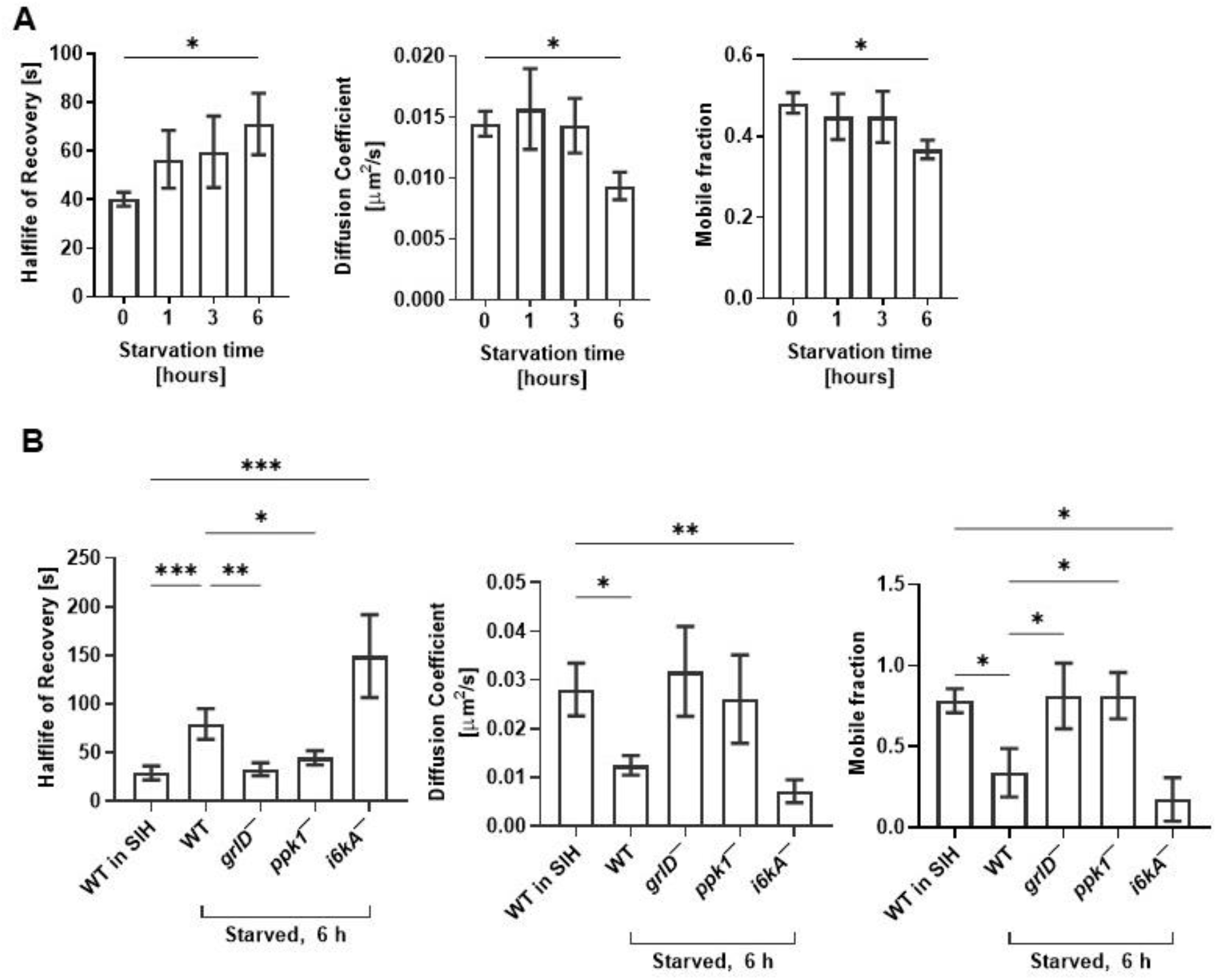
Starvation reduces cell membrane fluidity of WT and *i6kA*^−^ cells. **A-B)** Half-life of recovery, diffusion coefficient, and mobile fraction were calculated from fluorescence recovery after photobleaching of WT cells starved in PBM for 0, 1, 3 and 6 hours (A) and WT cells incubated in SIH for 0 h or WT, *grlD*^−^, *ppk1*^−^, and *i6kA*^−^ cells starved in PBM for 6 hours (B). Values are mean ± SEM of 3 independent experiments. * p < 0.05, ** p < 0.01, *** p < 0.001 (One-way ANOVA with Brown-Forsythe and Welch test).

### Starving *D. discoideum* cells require GrlD, Ppk1, and I6kA to reduce macropinocytosis and nutrient retention

As previously observed [15, 28], starvation for 6 hours decreased macropinocytosis of TRITC-dextran in WT cells (**Fig. 4A**). Starvation increased macropinocytosis in *grlD*^−^ cells and had no significant effect on *ppk1*^−^ and *i6kA*^−^ cells (**Fig. 4A**). Starvation reduces exocytosis [15], and in agreement with this we observed that after ingesting TRITC-dextran, and subsequently allowing 1 hour for excretion of ingested TRITC-dextran, WT cells starved for 6 hours retained more TRITC-dextran than freshly starved cells (**Fig. 4B**). Starvation for 6 hours did not significantly affect the retention of TRITC-dextran by *grlD*^−^, *ppk1*^−^, or *i6kA*^−^ cells (**Fig. 4B**). These results indicate that at 6 hours after starvation, cells require GrlD, Ppk1, and I6kA to inhibit micropinocytosis, decrease exocytosis, and increase retention of materials ingested by micropinocytosis.

**Figure 4.**
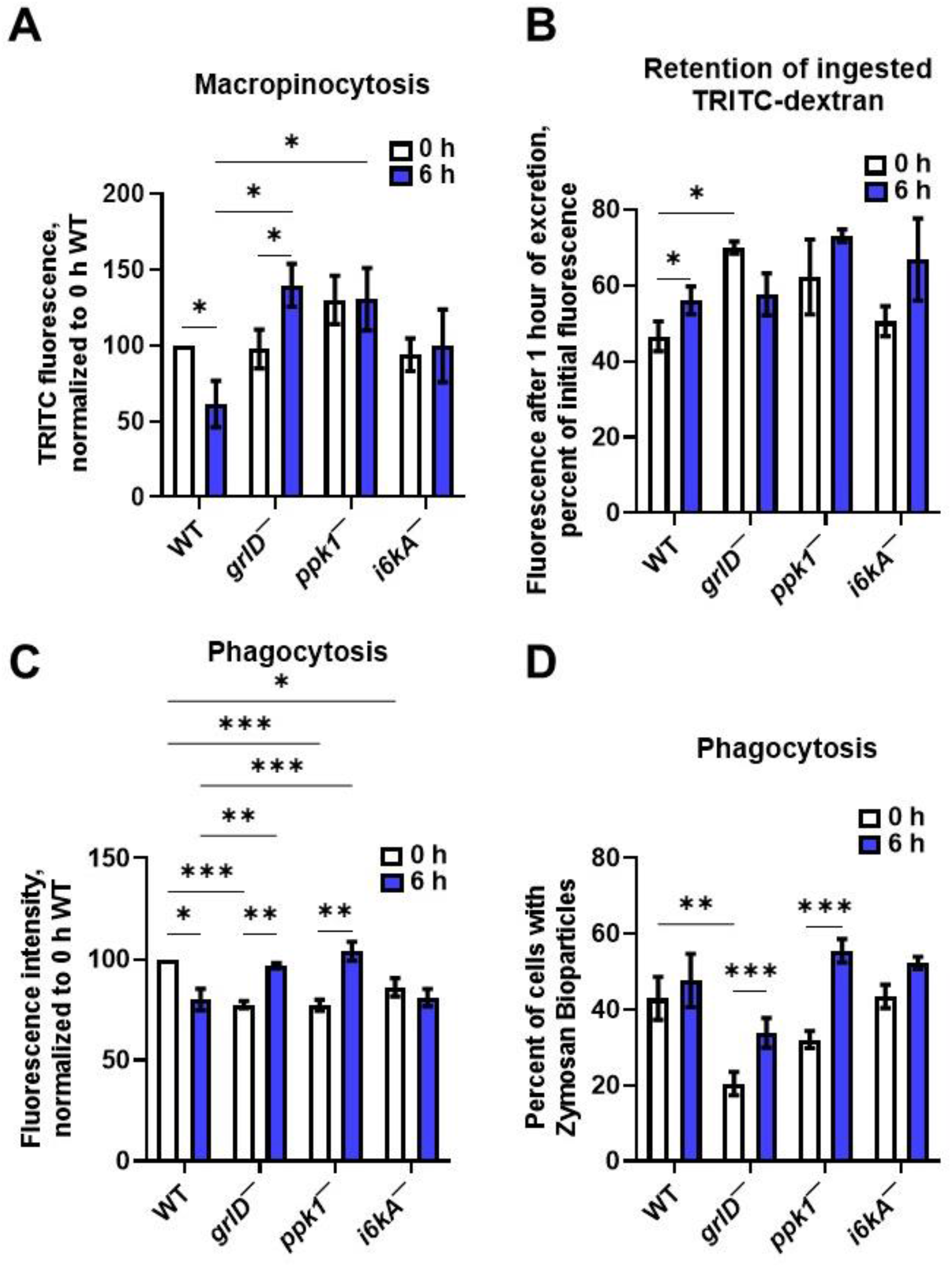
Starving *D. discoideum* cells require GrlD, Ppk1, and I6kA to reduce macropinocytosis, exocytosis, and phagocytosis. **A)** WT, *grlD*^−^, *ppk1*^−^, and *i6kA*^−^ cells starved in PBM for 0 or 6 hours were allowed to macropinocytose TRITC-dextran, and the median fluorescence intensity per cell of ingested TRITC-dextran were measured. The median fluorescence intensity per cell in WT cells starved for 0 h was considered 100%. **B)** WT, *grlD*^−^,*ppk1*^−^, and *i6kA*^−^ cells starved in PBM for 0 or 6 h were allowed to macropinocytose TRITC-dextran as in A, uningested TRITC-dextran was removed, and cells were allowed to excrete the ingested TRITC-dextran for 1 h. The median fluorescence intensities per cell of TRITC-dextran retained after 1 hour of excretion were determined. **C and D)** WT, *grlD*^−^, *ppk1*^−^, and *i6kA*^−^ cells were starved for 0 or 6 hours and allowed to phagocytose Zymosan-A bioparticles for 1 hour. The average median fluorescence intensity of ingested bioparticles, normalized to 0 h WT cells are shown in (C), and percentages of cells with ingested bioparticles are shown in (D). All values are mean ± SEM of 4 independent experiments. * indicates p < 0.05, ** p < 0.01, *** p < 0.001 (Two-way ANOVA with Šídák’s multiple comparisons test).

### Starving *D. discoideum* cells require GrlD, Ppk1, and I6kA to reduce phagocytosis

As with macropinocytosis (**Fig 4A**), and as previously observed [16], starvation for 6 hours reduced phagocytosis of yeast bioparticles in WT cells (**Fig. 4C**). This was not accompanied by a reduced percentage of cells with phagocytosed yeast (**Fig. 4D**). Starvation for 6 hours increased both phagocytosis and the percent of cells with phagocytosed yeast in *grlD^−^* and *ppk1*^−^ cells, but had no significant effect on these in *i6kA*^−^ cells (**Fig. 4C and D**).

### Cells lacking GrlD or Ppk1 are abnormally large

At 6 hours of starvation, *grlD*^−^ and *ppk1*^−^ cells had increased macropinocytosis and phagocytosis compared to WT cells, but no significant change in retention of ingested TRITC-dextran (**Fig. 4A-C**). Although during starvation there is probably little nutrients for cells to uptake by phagocytosis, the increased macropinocytosis of *grlD*^−^ and *ppk1*^−^ cells might increase their mass. To examine this, WT, *grlD*^−^, *ppk1*^−^, and *i6kA*^−^ were starved and cell size, cell mass, and total protein content were measured. Although *grlD*^−^ and *ppk1*^−^ cells tended to be larger and have more mass than WT cells (**Fig. 5A - C**), their mass did not significantly increase from 0 to 6 hours compared to WT cells, suggesting that the increased macropinocytosis of *grlD*^−^ and *ppk1*^−^ cells did not cause them to significantly gain additional mass over the first 6 hours of starvation. Compared to WT cells, *grlD*^−^ cells had more total protein content at all times examined, probably because they are intrinsically larger cells, and *ppk1*^−^ cells had more protein at 3 and 6 hours of starvation (**Fig. 5D**), suggesting that for unknown reasons they convert nutrient stores to protein.

**Figure 5.**
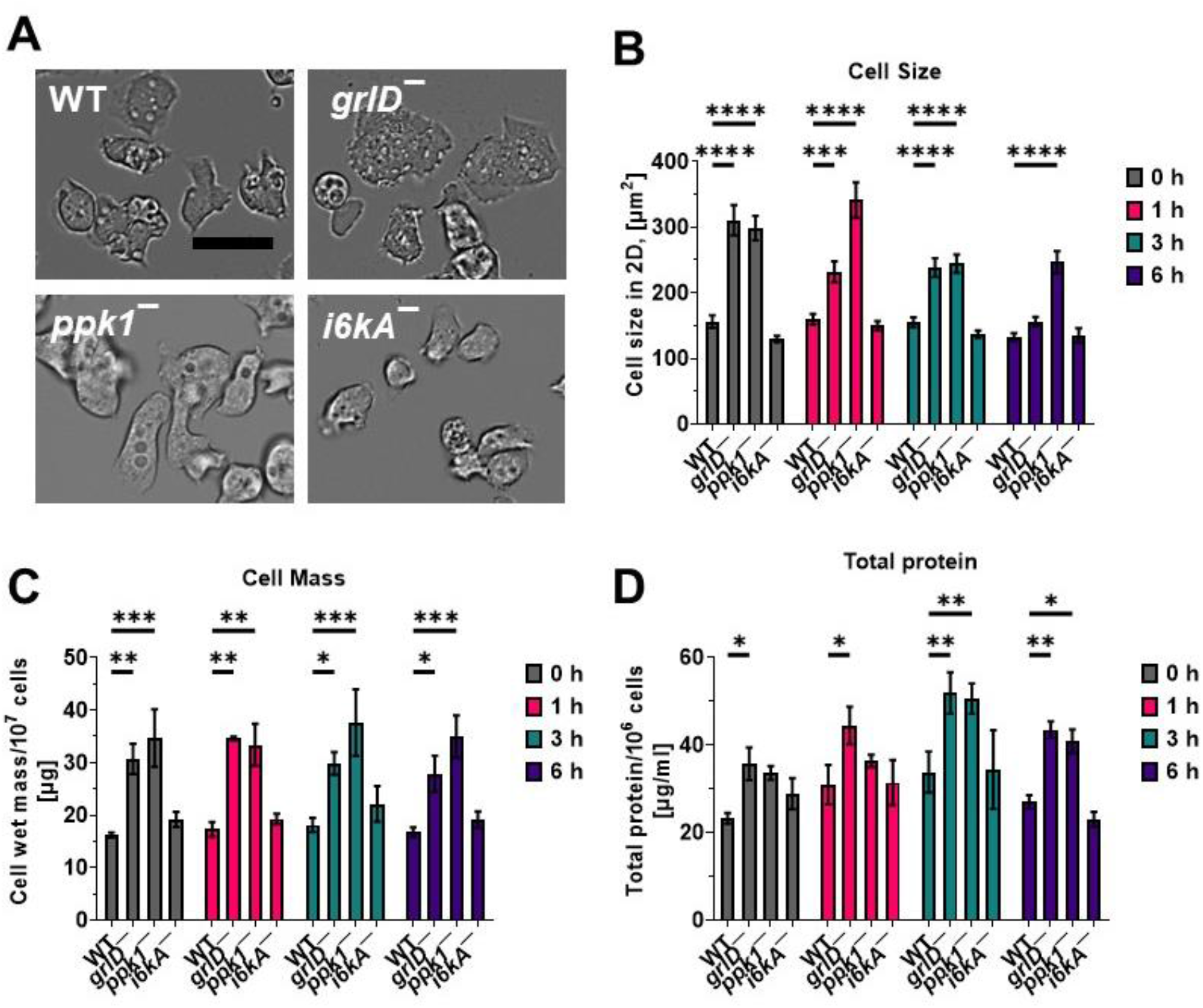
Starvation does not alter cell size, mass, and total protein content. **A)** Representative images of vegetative WT, *grlD*^−^, *ppk1*^−^, and *i6kA*^−^ cells growing in submerged stationary culture are shown from 3 independent experiments. Bar is 20 μm. **B)** Cell areas of 10^6^ cells/ml WT, *grlD*^−^, *ppk1*^−^, and *i6kA*^−^ cells starved in PBM in submerged stationary culture for 0, 1, 3, and 6 hours. At least 50 cells from three independent experiments were taken to measure the cell size. **C and D)** Cell mass per 10^7^ cells (C) and total protein content per 10^6^ cells (D) were determined for WT, *grlD*^−^, *ppk1*^−^, and *i6kA*^−^ cells starved in PBM for indicated times. All values are mean ± SEM of 3 independent experiments. For B-D, * indicates p < 0.05, ** p < 0.01, *** p < 0.001, and p < 0.0001 **** (Two-way ANOVA with Holm-Šídák’s multiple comparisons test).

## Discussion

PolyP is an autocrine signaling molecule in *D. discoideum*. Cells growing in the presence of nutrients (vegetative cells) secrete polyP, and at high cell densities where the cells are about to overgrow their food source, the concomitant high extracellular concentrations of polyP inhibit cell proliferation without compromising cell growth [27]. PolyP also primes cells to anticipate starvation and prepare them to enter starvation-induced development [24]. Here we showed that starved *D. discoideum* cells accumulate soluble and cell surface polyP and can bind exogenous polyP. PolyP inhibits the membrane fluidity of vegetative cells [28], and here we observed that compared to vegetative cells, starved cells have a decreased membrane fluidity.

Ppk1 synthesizes polyP, and as expected cells lacking Ppk1 have reduced accumulation of both extracellular and cell surface polyP during starvation. PolyP decreases membrane fluidity, and so also as expected, *ppk1*^−^ cells do not decrease membrane fluidity during starvation. Supporting the idea that polyP mediates the starvation-induced decrease in macropinocytosis, phagocytosis, and exocytosis, *ppk1*^−^ cells do not exhibit these decreases. Compared to WT cells, *ppk1*^−^ cells have reduced binding of exogenous polyP, suggesting the possibility that polyP increases levels of thereceptor that binds polyP.

GrlD is the receptor that binds and senses polyP, and as expected starved *grlD*^−^ cells show reduced binding of exogenous polyP. Unexpectedly, the *grlD*^−^ cells do not accumulate either extracellular or cell-surface polyP, suggesting that there exists some sort of feedback mechanism where sensing some polyP is needed to accumulate extracellular polyP. This might prevent cells from wasting energy releasing polyP in a situation where the cell is in an environment such as a rainstorm where the polyP will be immediately washed away. As a result of either not sensing or not accumulating extracellular polyP, cells lacking GrlD do not decrease membrane fluidity, macropinocytosis, phagocytosis, and exocytosis during starvation, further supporting the idea that sensing extracellular polyP mediates these decreases.

Along with Ppk1, I6kA is involved in the accumulation of polyP in vegetative cells, and in starved *i6kA*^−^ cells we observed normal levels of extracellular polyP but reduced levels of cell surface polyP. An intriguing possibility is that I6kA generates a polyP that has an inositol at one end, and that the cell surface polyP is associated with this hypothetical modified polyP. The *i6kA*^−^ cells show reduced binding of exogenous polyP, suggesting that I6kA affects the GrlD receptor directly, or that cell-surface polyP is needed for GrlD localization to the cell surface, or that cell-surface polyP is needed for GrlD function. Cells lacking I6kA show a normal decrease of membrane fluidity at 6 hours of starvation, but do not decrease macropinocytosis, phagocytosis, and exocytosis. This then indicates that during starvation, the increase in cell surface polyP is not responsible for the decrease in membrane fluidity, and that reduced cell surface polyP but not altered membrane fluidity is not responsible for the starvation effects on macropinocytosis, phagocytosis, and exocytosis.

High levels of polyP increase the size and mass of vegetative cells [27]. This would initially suggest that *grlD*^−^ cells that do not sense polyP [32], and *ppk1*^−^ cells, which make no detectable intracellular polyP [30] and have ~a 50% reduction of extracellular polyP [27], would tend to be small. However, we observed that vegetative and starved *grlD*^−^ and *ppk1*^−^ cells are abnormally large. For the *grlD*^−^ cells, one possible explanation is that in the apparent absence of any polyP, the cells sense that they have become isolated from a colony of other cells, and increase cell size and nutrient reserves in response to this abnormal situation. For the *ppk1*^−^ cells, one possible explanation is that making polyP is energetically costly, and not making intracellular polyP allows the cells to be both larger and proliferate faster than WT cells. Together these results suggest that in addition to acting as a cell density sensing signal during growth, polyPhas significant functions as a signal during development.

## Acknowledgements

The use of the Microscopy and Imaging Center facility at Texas A&M University is acknowledged. The Olympus FV1000 confocal microscope acquisition was supported by the Office of the Vice President for Research at Texas A&M University. Use of the TAMU/ Laboratory for Biological Mass Spectrometry service and collaboration facility (LBMS) is acknowledged. This work was supported by National Institutes of Health grants GM118355 and GM139486.

## Author Contributions

R. R. designed, performed experiments, analyzed data, and wrote the paper. I. I. and S. J. performed experiments, and R. H. G. coordinated the study and wrote the paper.

## Methods

### D. discoideum cell culture

WT AX2 (DBS0237699) [4], *grlD*^−^ (DBS0350227) [38], *ppk1*^−^ (DBS0350686) [30], and *i6kA*^−^ (DBS0236426) [39] *D. discoideum* strains were obtained from the *Dictyostelium* Stock Center [40]. *D. discoideum* cell cultures were maintained at 21 °C in type 353003 tissue culture dishes (Corning, Durham, NC) in 10 ml of SIH defined minimal medium (Formedium, Norfolk, England) under no selection (AX2) or selection with 5 μg/ml blasticidin (*grlD*^−^,*ppkl*^−^, and *i6kA*^−^). Cells were also grown on SM/5 agar [41] on lawns of *E. coli* DB (DBS0350636) in a type 2538-4302 petri dish (VWR, Radnor, PA). 100 μg/ml dihydrostreptomycin and 100 μg/ml ampicillin were used to kill *E. coli* in *D. discoideum* cultures obtained from SM/5 agar [42]. *D. discoideum* cells from 80% - 90% confluent cultures in a tissue culture dish were collected using a sterile glass pipette, transferred to 15 ml conical tubes (Falcon, VWR), washed 2 times with SIH by centrifugation at 500 x g for 5 minutes, the cell density was measured with a hemocytometer, and 300 μl of cells at 1 x 10^6^ cells/ml was transferred to the well of a type 353219 96-well, black/clear, tissue culture treated plate (Corning) to obtain 3×10^5^ cells per well, or 1 ml was transferred to the well of a type 353047 24-well tissue culture plate (Corning) to obtain 10^6^ cells per well. For starvation assays, *D. discoideum* cells in a 96-well, black/clear, tissue culture treated plate or a 24-well tissue culture plate were washed twice with PBM (20 mM KH2PO4, 0.01 mM CaCl2, and 1 mM MgCl2, pH adjusted to 6.1 with KOH) by centrifuging the plate at 500 x g for 3 minutes and replacing the supernatant with fresh PBM, and cells were incubated for 0, 1, 3, and 6 hours for starvation. The 24 and 96 well plates with cells were incubated in a Tupperware container with wet paper towels for humidity.

### Recombinant polyphosphatase purification and polyP concentration measurement

Recombinant *Saccharomyces cerevisiae* exopolyphosphatase (PPX) [35, 43] was purified following the method described for the purification of recombinant autocrine proliferation repressor protein AprA [44]. *D. discoideum* cells and culture supernatants were treated with PPX as previously described (Rijal et al., 2020). PolyP concentrations were determined from culture supernatants as previously described [28].

### PolyP binding assay and SYTOX staining of membrane-bound polyP

PolyP binding assays were performed as previously described [32], except that tag-free polyP was utilized instead of biotinylated polyP, and the bound polyP was stained with the polyP-binding fluorescent dye SYTOX as previously described [34]. Briefly, *D. discoideum* cells in a 96-well, black/clear, tissue culture-treated plate were spun down at 500 × g for 3 minutes. SIH medium was replaced with fresh SIH medium containing 705 μg/ml polyP and incubated for 3 minutes. Cells were spun down at 500 × g for 3 minutes and the medium was replaced with fresh SIH. This step was repeated twice to remove unbound polyP. SIH medium containing 1.5 μM SYTOX (Cat#S7020, Thermo Fisher Scientific, Waltham, MA) was added to the cells and incubated for 10 minutes, and images were taken using a 40× objective on a Nikon Eclipse Ti2 inverted microscope (Nikon, Melville, NY). Deconvolution of images was done using the Richardson-Lucy algorithm [45] in Nikon NIS-Elements AR software. The fluorescence intensity of SYTOX staining was analyzed by ImageJ [46]. SYTOX staining of cells in SIH or cells starved in PBM for 2, 3, or 6 hours was performed as described above but in the absence of added exogenous polyP. Where indicated, 5 μg/ml PPX was added to the cells incubated in the PBM during the 2, 3, or 6 hours of starvation.

### DNase and RNase treatments

Cells were starved in PBM for 5.5 hours in a 96-well, black/clear, tissue culture-treated plate as described above and 50 μg/ml RNase (Cat#109142, Roche CustomBiotech, Indianapolis, IN) or 50 μg/ml DNase (04536282001, Roche Diagnostics, Mannheim, Germany) were added to cells, and the cells were incubated for 30 minutes, SYTOX staining of the cells was performed, and images of the cells were taken as described above. To test the enzymatic activity of RNase and DNase, 1 μg of total RNA isolated from WT *D. discoideum* cells as previously described [47] or 1 μg of a plasmid DNA PDM232 [48] was incubated without or with 50 μg/ml of RNase or DNase, respectively, for 45 minutes at room temperature, and resolved by 0.7% agarose gel electrophoresis with ethidium bromide stain [49].

### Fluorescence recovery after photobleaching

Photobleaching assays were done as previously described [28, 50], except that the *D. discoideum* cells were starved in PBM buffer for the indicated times.

### Macropinocytosis and exocytosis assays

For both macropinocytosis and exocytosis, tetramethylrhodamine isothiocyanate-dextran (TRITC-dextran) (T1162-100MG, Sigma-Aldrich, St. Louis, MO) was used to visualize ingestion and retention [28, 51]. For macropinocytosis, cells were starved in a 24-well plate in PBM with 5 μl of 1 mg/ml TRITC-dextran for 1 hour to allow macropinocytosis of the TRITC-dextran. After 1 hour, the cells incubated with TRITC-dextran were spun down at 500 x g for 3 minutes, the supernatant was replaced with fresh PBM, and this step was repeated twice. Cells were gently washed off the bottom of the plate with 200 μl PBM. The median fluorescence of the live cell population was recorded using the PE-A fluorescence gate on a Accuri C6 flow cytometer (BD, San Jose, CA). The same procedure was repeated for the 6 hour starved cells by adding TRITC-dextran at 6 hours of starvation and collecting the cells at 7 hours.

The exocytosis assay was performed similarly to the endocytosis assay. Freshly starved cells were incubated with TRITC-dextran for 1 hour, and the cells were washed 3 times with PBM as above, an aliquot of cells was collected and the fluorescence of the retained ingested TRITC-dextran was measured using the flow cytometer, and 1 ml of PBM was then added to the remaining cells to allow exocytosis. After 1 hour, cells were collected, and the fluorescence of the retained ingested TRITC-dextran was measured. Similarly, 6-hour starved cells were incubated with TRITC-dextran for 1 hour, and the cells were washed as above with PBM, and 1 ml of conditioned medium collected from cells starved for 6 hours in PBM in a separate culture were added to the cells with ingested TRITC-dextran to allow exocytosis for 1 hour. The percentage of TRITC-dextran retained after exocytosis was calculated from the median fluorescence of the ingested TRITC-dextran in the cells that were allowed to exocytose for 1 hour divided by the median fluorescence of the ingested TRITC-dextran in the cells that were not allowed to exocytose.

### Phagocytosis assay

*D. discoideum* cells were starved for 0 and 6 hours as described above. The cells were collected and 800 μl was used to measure background fluorescence intensity in the PerCP-A channel in the flow cytometer. The remaining 200 μl in the wells were incubated with 5 μl of 1 mg/ml of Zymosan-A yeast BioParticles (Cat#Z23374, Thermo Fisher Scientific) in PBM and allowed to phagocytose for 1 hour. Cells were collected and the median fluorescence of the live cell population was measured using the fluorescence gating for PerCP-A on the flow cytometer.

### Cell size, mass, and protein determination

To determine cell size, mass, and total protein content, 50 ml of cells at 1 x 10^6^ cells/ml were starved as described above in a shaking culture for 0, 1, 3, or 6 hours. For cell size measurement, at each time point, 100 μl of cells were transferred to a 96-well, black/clear, tissue culture-treated plate, and images of cells were taken using a 40× objective on a Nikon Eclipse Ti2 inverted microscope. Cell sizes were measured using Fiji (ImageJ; NIH). At least 50 cells from each of three independent experiments were used to measure cell size.

For cell mass and total protein content measurement, cells from 10 ml of starved cells were collected by centrifugation at 500 x g for 3 minutes. The pellet was resuspended in approximately 500 μl of residual PBM and transferred to microcentrifuge tubes. The cells were collected by centrifugation at 3000 x g for 3 minutes. The supernatant was discarded, and the cell pellets were weighed. The total protein content of cells was determined as previously described [52].

### Statistical analysis

Statistical analysis was performed using Prism 9 (GraphPad, San Diego, CA) and the tests indicated in the figure legends. A p < 0.05 was considered to be significant.

## References

1. Cardelli, J. “Phagocytosis and Macropinocytosis in Dictyostelium: Phosphoinositide-Based Processes, Biochemically Distinct.” Traffic 2, no. 5 (2001): 311–20.

2. Vuillemin, PAUL. “Une Acrasiee Bacteriophage.” Comptes rendus des s eances de I’Acad emie des sciences de Paris 137 (1903): 387–89.

3. Cosson, P., and T. Soldati. “Eat, Kill or Die: When Amoeba Meets Bacteria.” Curr Opin Microbiol 11, no. 3 (2008): 271–6.

4. Watts, DJ, and JM Ashworth. “Growth of Myxamoebae of the Cellular Slime Mould Dictyostelium Discoideum in Axenic Culture.” Biochemical Journal 119, no. 2 (1970): 171.

5. Franke, J., and R. Kessin. “A Defined Minimal Medium for Axenic Strains of Dictyostelium Discoideum.” Proc Natl Acad Sci U S A 74, no. 5 (1977): 2157–61.

6. Watts, DJ. “Vitamin Requirements for Growth of Myxamoebae of Dictyostelium Discoideum in a Defined Medium.” Microbiology 98, no. 2 (1977): 355–61.

7. Sussman, R., and M. Sussman. “Cultivation of Dictyostelium Discoideum in Axenic Medium.” Biochem Biophys Res Commun 29, no. 1 (1967): 53–5.

8. Hacker, U., R. Albrecht, and M. Maniak. “Fluid-Phase Uptake by Macropinocytosis in Dictyostelium.” J Cell Sci 110 (Pt 2), no. 2 (1997): 105–12.

9. Loomis, W. F., Jr. “Sensitivity of Dictyostelium Discoideum to Nucleic Acid Analogues.” Exp Cell Res 64, no. 2 (1971): 484–6.

10. Bloomfield, G., D. Traynor, S. P. Sander, D. M. Veltman, J. A. Pachebat, and R. R. Kay. “Neurofibromin Controls Macropinocytosis and Phagocytosis in Dictyostelium.” eLife 4 (2015): e04940.

11. Maniak, M. “Fusion and Fission Events in the Endocytic Pathway of Dictyostelium.” Traffic 4, no. 1 (2003): 1–5.

12. Neuhaus, E. M., W. Almers, and T. Soldati. “Morphology and Dynamics of the Endocytic Pathway in Dictyostelium Discoideum.” Mol Biol Cell 13, no. 4 (2002): 1390–407.

13. Raper, Kenneth B. “Dictyostelium Discoideum, a New Species Of.” Journal of agricultural research 50 (1935): 135.

14. Kessin, Richard H. Dictyostelium: Evolution, Cell Biology, and the Development of Multicellularity. Vol. 38: Cambridge University Press, 2001.

15. Smith, E. W., W. C. Lima, S. J. Charette, and P. Cosson. “Effect of Starvation on the Endocytic Pathway in Dictyostelium Cells.” Eukaryot Cell 9, no. 3 (2010): 387–92.

16. Katoh, M., G. Chen, E. Roberge, G. Shaulsky, and A. Kuspa. “Developmental Commitment in Dictyostelium Discoideum.” Eukaryot Cell 6, no. 11 (2007): 2038–45.

17. Zhang, X., O. Zhuchenko, A. Kuspa, and T. Soldati. “Social Amoebae Trap and Kill Bacteria by Casting DNA Nets.” Nat Commun 7, no. 1 (2016): 10938.

18. Chen, G., O. Zhuchenko, and A. Kuspa. “Immune-Like Phagocyte Activity in the Social Amoeba.” Science 317, no. 5838 (2007): 678–81.

19. Rao, N. N., M. R. Gomez-Garcia, and A. Kornberg. “Inorganic Polyphosphate: Essential for Growth and Survival.” Annu Rev Biochem 78 (2009): 605–47.

20. Kornberg, Arthur, Narayana N Rao, and Dana Ault-Riche. “Inorganic Polyphosphate: A Molecule of Many Functions.” Annual review of biochemistry 68, no. 1 (1999): 89–125.

21. Crooke, E., M. Akiyama, N. N. Rao, and A. Kornberg. “Genetically Altered Levels of Inorganic Polyphosphate in Escherichia Coli.” J Biol Chem 269, no. 9 (1994): 6290–5.

22. Rashid, M Harunur, Kendra Rumbaugh, Luciano Passador, David G Davies, Abdul N Hamood, Barbara H Iglewski, and Arthur Kornberg. “Polyphosphate Kinase Is Essential for Biofilm Development, Quorum Sensing, and Virulence of Pseudomonas Aeruginosa.” Proceedings of the National Academy of Sciences 97, no. 17 (2000): 9636–41.

23. Schroder, H. C., L. Kurz, W. E. Muller, and B. Lorenz. “Polyphosphate in Bone.” Biochemistry (Mosc) 65, no. 3 (2000): 296–303.

24. Suess, P. M., J. Watson, W. Chen, and R. H. Gomer. “Extracellular Polyphosphate Signals through Ras and Akt to Prime Dictyostelium Discoideum Cells for Development.” J Cell Sci 130, no. 14 (2017): 2394–404.

25. Hassanian, S. M., A. Avan, and A. Ardeshirylajimi. “Inorganic Polyphosphate: A Key Modulator of Inflammation.” J Thromb Haemost 15, no. 2 (2017): 213–18.

26. Smith, S. A., N. J. Mutch, D. Baskar, P. Rohloff, R. Docampo, and J. H. Morrissey. “Polyphosphate Modulates Blood Coagulation and Fibrinolysis.” Proc Natl Acad Sci U S A 103, no. 4 (2006): 903–8.

27. Suess, P. M., and R. H. Gomer. “Extracellular Polyphosphate Inhibits Proliferation in an Autocrine Negative Feedback Loop in Dictyostelium Discoideum.” J Biol Chem 291, no. 38 (2016): 20260–9.

28. Rijal, R., S. A. Kirolos, R. J. Rahman, and R. H. Gomer. “Dictyostelium Discoideum Cells Retain Nutrients When the Cells Are About to Outgrow Their Food Source.” J Cell Sci 135, no. 18 (2022).

29. Zhang, H., M. R. Gomez-Garcia, X. Shi, N. N. Rao, and A. Kornberg. “Polyphosphate Kinase 1, a Conserved Bacterial Enzyme, in a Eukaryote, Dictyostelium Discoideum, with a Role in Cytokinesis.” Proc Natl Acad Sci U S A 104, no. 42 (2007): 16486–91.

30. Livermore, T. M., J. R. Chubb, and A. Saiardi. “Developmental Accumulation of Inorganic Polyphosphate Affects Germination and Energetic Metabolism in Dictyostelium Discoideum.” Proc Natl Acad Sci U S A 113, no. 4 (2016): 996–1001.

31. Saiardi, A., H. Erdjument-Bromage, A. M. Snowman, P. Tempst, and S. H. Snyder. “Synthesis of Diphosphoinositol Pentakisphosphate by a Newly Identified Family of Higher Inositol Polyphosphate Kinases.” Curr Biol 9, no. 22 (1999): 1323–6.

32. Suess, P. M., Y. Tang, and R. H. Gomer. “The Putative G Protein-Coupled Receptor Grld Mediates Extracellular Polyphosphate Sensing in Dictyostelium Discoideum.” Mol Biol Cell 30, no. 9 (2019): 1118–28.

33. Klein, Gerard, David A Cotter, Jean Baptiste Martin, Mireille Bof, and Michel Satre. “Germination of Dictyostelium Discoideum Spores. A Phosphorus-31 Nmr Analysis.” Biochemistry 27, no. 21 (1988): 8199–203.

34. Verhoef, J. J., A. D. Barendrecht, K. F. Nickel, K. Dijkxhoorn, E. Kenne, L. Labberton, O. J. McCarty, R. Schiffelers, H. F. Heijnen, A. P. Hendrickx, H. Schellekens, M. H. Fens, S. de Maat, T. Renne, and C. Maas. “Polyphosphate Nanoparticles on the Platelet Surface Trigger Contact System Activation.” Blood 129, no. 12 (2017): 1707–17.

35. Wurst, H., and A. Kornberg. “A Soluble Exopolyphosphatase of Saccharomyces Cerevisiae. Purification and Characterization.” J Biol Chem 269, no. 15 (1994): 1099–61001.

36. Ben-Dov, N., and R. Korenstein. “Proton-Induced Endocytosis Is Dependent on Cell Membrane Fluidity, Lipid-Phase Order and the Membrane Resting Potential.” Biochim Biophys Acta 1828, no. 11 (2013): 2672–81.

37. Ge, S., J. G. White, and C. L. Haynes. “Critical Role of Membrane Cholesterol in Exocytosis Revealed by Single Platelet Study.” ACS Chem Biol 5, no. 9 (2010): 819–28.

38. Tang, Y., Y. Wu, S. E. Herlihy, F. J. Brito-Aleman, J. H. Ting, C. Janetopoulos, and R. H. Gomer. “An Autocrine Proliferation Repressor Regulates Dictyostelium Discoideum Proliferation and Chemorepulsion Using the G Protein-Coupled Receptor Grlh.” mBio 9, no. 1 (2018).

39. Luo, H. R., Y. E. Huang, J. C. Chen, A. Saiardi, M. Iijima, K. Ye, Y. Huang, E. Nagata, P. Devreotes, and S. H. Snyder. “Inositol Pyrophosphates Mediate Chemotaxis in Dictyostelium Via Pleckstrin Homology Domain-Ptdins(3,4,5)P3 Interactions.” Cell 114, no. 5 (2003): 559–72.

40. Fey, Petra, Robert J. Dodson, Siddhartha Basu, and Rex L. Chisholm. “One Stop Shop for Everything Dictyostelium: Dictybase and the Dicty Stock Center in 2012.” In Dictyostelium Discoideum Protocols, edited by Ludwig Eichinger and Francisco Rivero, 59–92. Totowa, NJ: Humana Press, 2013.

41. Sussman, Maurice. “Chapter 14 Biochemical and Genetic Methods in the Study of Cellular Slime Mold Development.” In Methods in Cell Biology, edited by David M. Prescott, 397–410: Academic Press, 1966.

42. Brock, D. A., and R. H. Gomer. “A Cell-Counting Factor Regulating Structure Size in Dictyostelium.” Genes Dev 13, no. 15 (1999): 1960–9.

43. Gray, M. J., W. Y. Wholey, N. O. Wagner, C. M. Cremers, A. Mueller-Schickert, N. T. Hock, A. G. Krieger, E. M. Smith, R. A. Bender, J. C. Bardwell, and U. Jakob. “Polyphosphate Is a Primordial Chaperone.” Mol Cell 53, no. 5 (2014): 689–99.

44. Brock, D. A., and R. H. Gomer. “A Secreted Factor Represses Cell Proliferation in Dictyostelium.” Development 132, no. 20 (2005): 4553–62.

45. Laasmaa, M., M. Vendelin, and P. Peterson. “Application of Regularized Richardson-Lucy Algorithm for Deconvolution of Confocal Microscopy Images.” J Microsc 243, no. 2 (2011): 124–40.

46. Shihan, M. H., S. G. Novo, S. J. Le Marchand, Y. Wang, and M. K. Duncan. “A Simple Method for Quantitating Confocal Fluorescent Images.” Biochem Biophys Rep 25 (2021): 100916.

47. Rijal, R., K. M. Consalvo, C. K. Lindsey, and R. H. Gomer. “An Endogenous Chemorepellent Directs Cell Movement by Inhibiting Pseudopods at One Side of Cells.” Mol Biol Cell 30, no. 2 (2019): 242–55.

48. Veltman, D. M., G. Akar, L. Bosgraaf, and P. J. Van Haastert. “A New Set of Small, Extrachromosomal Expression Vectors for Dictyostelium Discoideum.” Plasmid 61, no. 2 (2009): 110–8.

49. Aaij, C., and P. Borst. “The Gel Electrophoresis of DNA.” Biochim Biophys Acta 269, no. 2 (1972): 192–200.

50. Tanaka, M., T. Kikuchi, H. Uno, K. Okita, T. Kitanishi-Yumura, and S. Yumura. “Turnover and Flow of the Cell Membrane for Cell Migration.” Sci Rep 7, no. 1 (2017): 12970.

51. Rivero, Francisco, and Markus Maniak. “Quantitative and Microscopic Methods for Studying the Endocytic Pathway.” In Dictyostelium Discoideum Protocols, edited by Ludwig Eichinger and Francisco Rivero, 423–38. Totowa, NJ: Humana Press, 2006.

52. Phillips, J. E., and R. H. Gomer. “The Roco Kinase Qkga Is Necessary for Proliferation Inhibition by Autocrine Signals in Dictyostelium Discoideum.” Eukaryot Cell 9, no. 10 (2010): 1557–65.

